# Tormentor: An obelisk prediction and annotation pipeline

**DOI:** 10.1101/2024.05.30.596730

**Authors:** Frederico Schmitt Kremer, Danielle Ribeiro de Barros

## Abstract

Obelisks are circular RNA viroid-like elements first described in January 2024 and identified through RNA-Seq data analysis. These elements are widespread in human gut samples. and characterized by their rod-like secondary structure reminiscent of obelisk monuments, exhibiting an average size of approximately 1,000 base pairs. Despite their initial identification, little is known about their biogenesis and their potential impact on human physiology and the microbiome. To facilitate further exploration in this emerging field, we developed Tormentor, an automated pipeline designed for the identification and annotation of Obelisks from stranded RNA-Seq data, leveraging methodologies similar to those employed in their initial discovery. To our knowledge, this is the first tool specialized for the identification of these elements.

## Full Text

Viroids are circular RNA elements that differ from viruses mostly due to their lack of protein coating (Tsagris *et al*. 2008). Although many known viroids are phytopathogens, the application of metatranscriptome has demonstrated that these elements, and similar ones (viroid-like), are widely present and distributed on RNA-Seq data froma wide variety of organisms (Lee *et al*. 2023).

Obelisks are circular RNA viroid-like elements described for the first time in January 2024 from RNA-Seq data, with an average size of ∼ 1.000 base pairs (Zheludev *et al*. 2024). The name refers to their secondary structure, mostly composed of self-pairing regions that form an “obelisk” (rod-like) shape. They usually present ORFs that potentially encode for “oblin” proteins which are conserved among different obelisks. Two main oblins are known: oblin-1 (a hypothetical globular protein) and oblin-2 (a hypothetical leucine zipper alpha helix protein). However, due to the lack of homology of these proteins to other sequences in protein databases, the function and structure are unknown and the only information available is provided by prediction methods. The original pre-print describing their existence and structure demonstrated that they are widely distributed on the human gut and in other NGS datasets as well, although little information is available about their biogenesis and effect on the human body. Additionally, another work, by Maddamsetti & You (2024) demonstrated that these elements are widely present in the transcriptome of *Streptococcus sanguinis* SK36, but not in its genome.

To facilitate further research on the new class of viroid-like elements, we have developed an automated pipeline named Tormentor, which performs the identification and annotation of Obelisks from stranded RNA-Seq data, following most of the steps used Zheludev *et al* (2024) on the first report. To our knowledge, this is the first automated pipeline available for the prediction of this specific kind of element.

The pipeline was implemented using the Python programming language version 3.8 (https://www.python.org/). Raw RNA-Seq reads are processed and trimmed using FASTP (Chen *et al*. 2018) and assembled using rnaSPAdes (Bushmanova *et al*. 2019), followed by an identification of viroid-like elements using VNOM (https://github.com/Zheludev/VNom). Circular elements with lengths between 700 and 2000 (by default) are kept and analyzed using Prodigal (Hyatt *et al*. 2010) for ORF annotation and adjustment of the sequence phase based on the start position of the largest ORF. Obelisks-related ribozyme motifs are annotated using `cmscan` from the INFERNAL (Nawrocki and Eddy 2013) package using the covariance models developed by Zheludev *et al* (2024), and secondary structures and predicted using RNAFold (Gruber *et al*. 2008). Only sequences containing ORFs with at least one hit (70% of similarity) on BLAST (Camacho *et al*. 2009) to the sequence of *oblin* proteins and a secondary structure composed of at least 70% of self-pairing regions are considered potential obelisks.

To evaluate the proposed pipeline we have analyzed the metatranscriptome of the SRA study SRR5949245, derived from the Human Microbiome Project. Raw data was obtained from NCBI SRA using the `fasterq-dump` tool from the SRA-Toolkit (https://github.com/ncbi/sra-tools) and processed on an Ubuntu Linux machine.

The pipeline is freely available on GitHub (https://github.com/omixlab/tormentor) and the main dependencies can be installed using conda (https://docs.anaconda.com/free/miniconda/index.html) and Make (https://www.gnu.org/software/make/). As input, the user must only provide a dataset of stranded paired-end reads in FASTQ, an output directory, and the path to the dataset directory (the directory is present in the GitHub repository). Additionally, it is also possible to define the minimum percent of self-paring in the obelisk candidates, by default set to “0.7”, e-value and identity thresholds for BLAST, number of threads to be used by rnaSPAdes, along with other optional arguments.

Using our pipeline to analyze the SRA study SRR5949245, we have identified an obelisk structure that matches the description made by Zheludev *et al*. (2024). Two oblin-coding ORF were identified (oblin-1 and oblin-2), along with obelisk-related ribozyme sites, a high percent of self-pairing bases. Finally, the circular plot representing the annotation of the predicted obelisk is presented in Figure 2.A, and the rod-like secondary structure predicted by RNAfold is presented in Figure 2.B.

**Figure 1.**
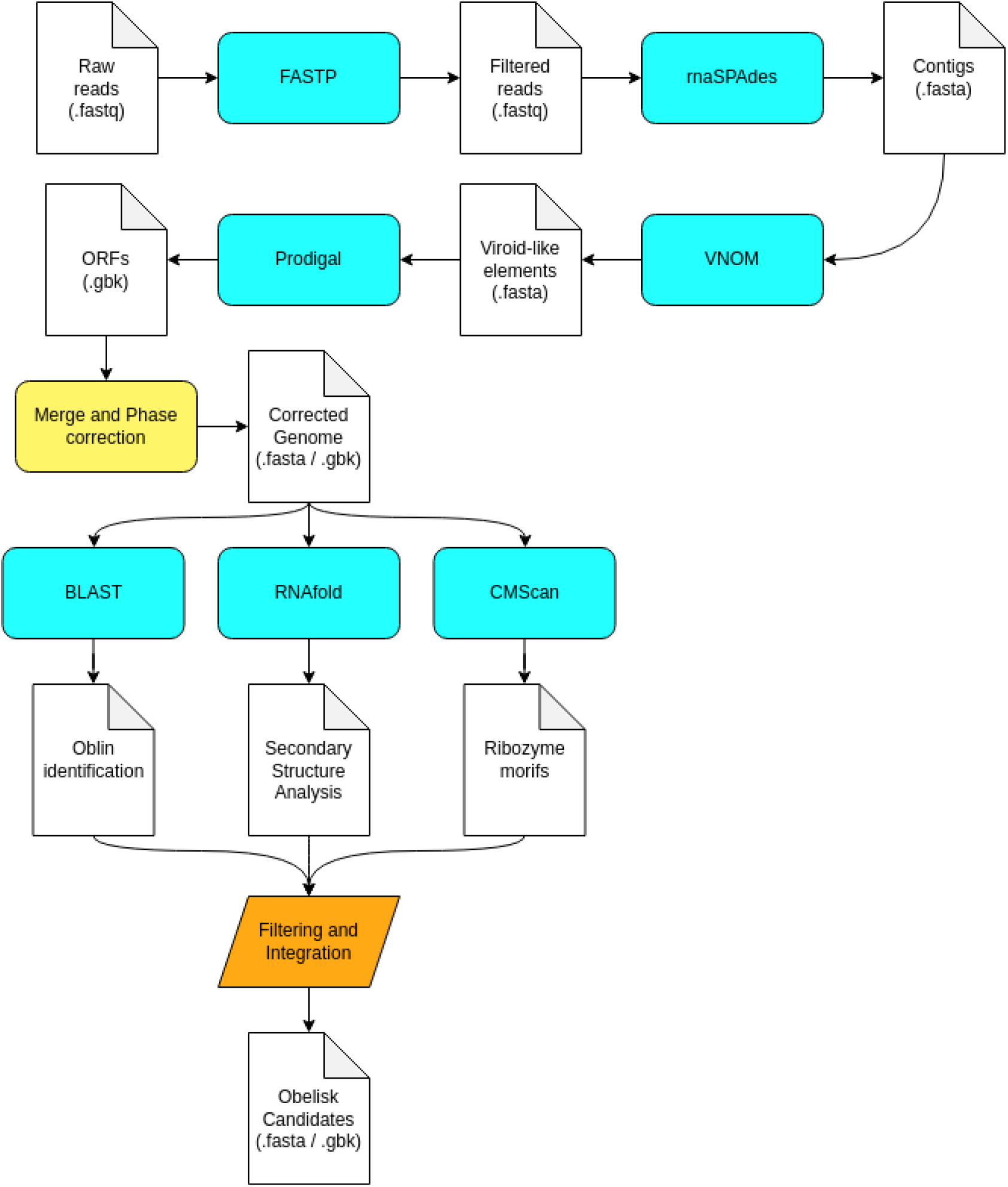
A workflow representing the Tormentor pipeline. Raw reads are processed using FASP and assembled into transcript contigs using rnaSPAdes. VNOM is used to identify circular transcripts and viroid-like elements. Prodigal is used to identify Open Reading Frames (ORFs) on the viroid-like elements and its results are used to adjust the phase of the sequence. BLAST is used to identify the oblin proteins, RNAfold is used to analyze the secondary structure, and cmscan, from the INFERNAL package, is used to identify ribozyme motifs characteristic of some Obelisks. All annotation results are merged and the final set of obelisks are saved in the output directory.

**Figure 2.**
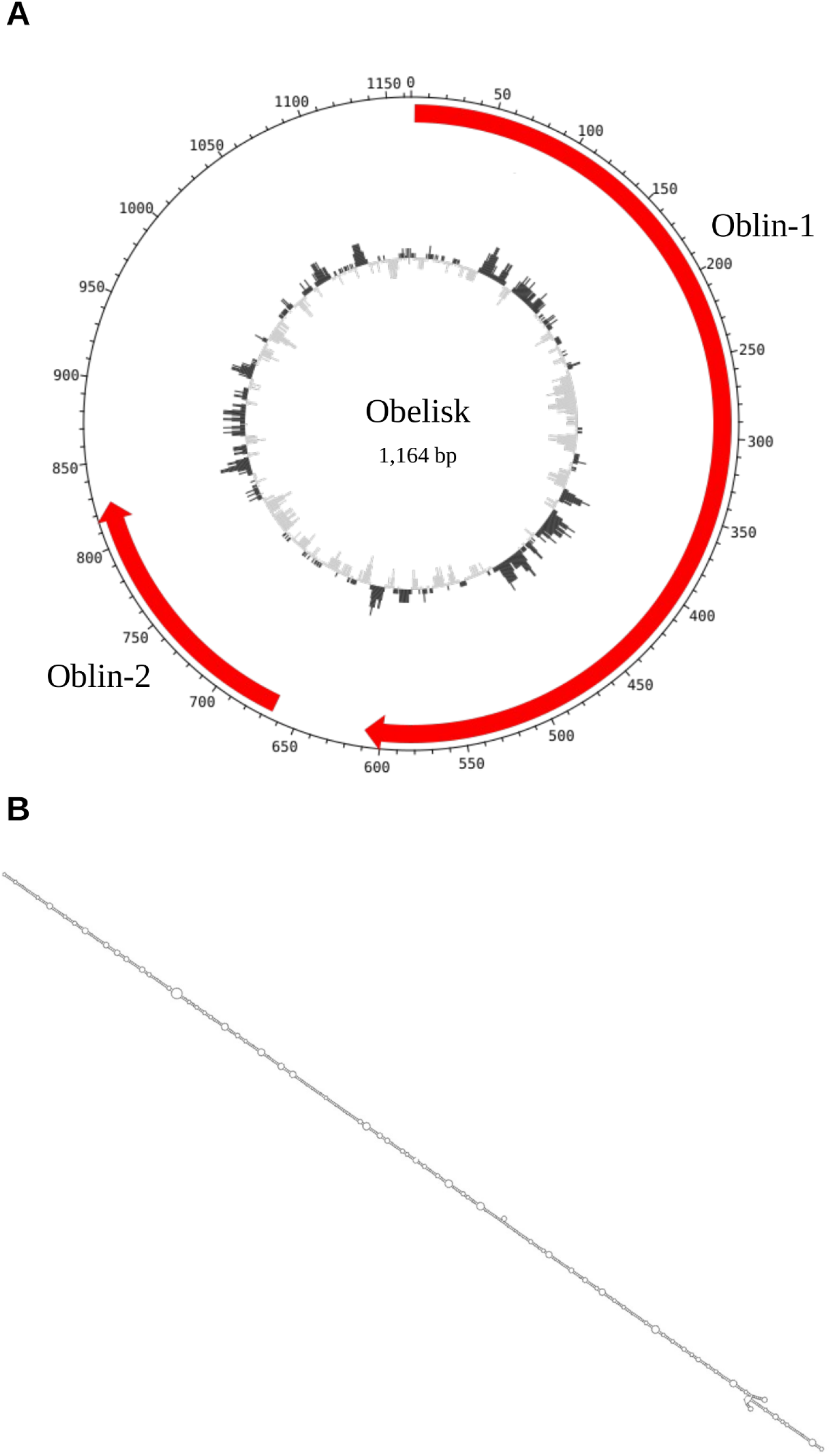
Obelisk identified in the SRA study SRR5949245 derived from the Human Microbiome Project (A) Annotation and circular genome plot. (B) The secondary structure of the obelisk illustrates the rod-like shape.

Metagenomics and metatranscriptomics have already contributed to a more complete understanding of the diversity and distribution of viruses and related elements (Rosario, Duffy and Breitbart 2012;Shi *et al*. 2016; Lee *et al*. 2023). Although still new in comparison to more classic techniques (eg: phenotypic analysis, serology), these approaches, when supported by consistent bioinformatics and genomic analysis, are already recognized by the International Committee on Taxonomy of Viruses (ICTV) for the taxonomic assignment of new species and other taxonomic ranks (Simmonds *et al*. 2017).

Although viroids are known since the decade 1970 (Diener 1979), the understanding of the distribution of viroid-like elements is still very limited, and only in the last years it was revealed that they are not only intracellular parasites of plants but also present in the human microbiome and the transcriptome of certain bacteria (Lee *et al*. 2023). The current understanding of Obelisks is even more limited, and only in early 2024, their first report was made, with a first proposal of characteristics that distinguish this class of elements from the other already characterized viroid-like elements. However, due to their novelty, to our knowledge, no proper tool is available to easily identify them. Therefore, with Tormentor, we hope to accelerate and facilitate the identification of these elements, which might help in the further understanding of their evolution, host interactions, and distribution.

Furthermore, the widespread distribution of obelisks across diverse datasets, including microbial species like *Streptococcus sanguinis*, a normal constituent of the human mouth microbiome but that can also be pathogenic when in the bloodstream, which leads to systemic disease (Nobbs and Kreth 2019), underscores their potential ecological significance. Additionally, their presence in microbial transcriptomes suggests potential interactions with host organisms or environmental factors, warranting exploration into their ecological roles and evolutionary dynamics.

Looking ahead, future research efforts should focus on refining our understanding of obelisks’ biological function and ecological relevance. Improved computational tools, like Tormentor, will be instrumental in advancing obelisk research, enabling researchers to efficiently analyze large-scale transcriptomic datasets and uncover novel insights into these intriguing viroid-like circular RNA elements.

Tormentor has shown to be useful in the identification of the novelly discovered class of viroid-like elements called Obelisks, and present results consistent with the reports that described these elements originally. However, the pipeline must still be improved, and future new features include machine-learning-based filtering of Obelisks and classification into sub-groups using kmer content, and a more extensive analysis of the secondary structure of the RNA to better identify the rod-like characteristic folding. The pipeline is freely available at GitHub (https://github.com/omixlab/tormentor) and might accelerate the identification of annotations of this newly discovered viroid-like element.

